# Antigen-scaffolds loaded with hyper-stable Neoleukin-2/15 expand antigen-specific T cells with a favorable phenotype for adoptive cell therapy

**DOI:** 10.1101/2025.04.16.649086

**Authors:** Maria Ormhøj, Kamilla Kjærgaard Munk, Siri Tvingsholm, Keerthana Ramanathan, Amalie Kai Bentzen, Georgios Kladis, Gitte N. Aasbjerg, Grigorii Nos, Tripti Tamhane, Hólmfridur Rósa Halldórsdóttir, Søren Nybo, Marcus Svensson Frej, Kristoffer Haurum Johansen, Mohammad Kadivar, Sine Reker Hadrup

## Abstract

Adoptive cell therapy (ACT) has shown promising results in cancer treatment, however, achieving effective *ex vivo* expansion of potent, functionally active, and cytotoxic T cells remains challenging. To overcome this, we loaded the engineered cytokine Neoleukin-2/15 (Neo2/15) on our recently established artificial antigen-presenting scaffolds (Ag-scaffolds) to expand antigen-specific T cells. Neo2/15 selectively binds to IL-2Rβ/γ receptors, enhancing CD8^+^ T cell proliferation while limiting regulatory T cell expansion.

Our study assessed the efficacy of Neo2/15-loaded Ag-scaffolds (Ag-Neo2/15 scaffolds) in expanding antigen-specific T cells from peripheral blood mononuclear cells (PBMCs) of healthy donors. We optimized Ag-scaffold configurations by varying the number of Neo2/15 molecules loaded on Ag-scaffolds and evaluated their impact on T-cell expansion and functionality. We showed that Ag-Neo2/15 scaffolds promoted significant T-cell expansion, with a comparable frequency of antigen-specific CD8^+^ T cells compared to IL-2/IL-21-loaded Ag-scaffolds (Ag-IL2/21 scaffolds). The CD8^+^ T cells expanded with Ag-Neo2/15 scaffolds exhibited potent TNFα and IFNγ production and expressed high levels of α4β7 integrin, a homing molecule which is important for directing T cells to specific tissues, potentially enhancing their therapeutic potential. T cells expanded with Ag-Neo2/15 scaffolds had superior and durable cytotoxicity against tumor target cells compared to T cells expanded with Ag-IL2/21 scaffolds. These findings were further supported by our single-cell analysis revealing that T cells expanded with Ag-Neo2/15 scaffolds had higher cytotoxic scores and lower dysfunctionality scores compared to T cells expanded with Ag-IL2/21 scaffolds. The single-cell analysis also indicated increased expression of genes linked to cell division and enhanced proliferative capacity in Ag-Neo2/15 expanded T cells. Furthermore, TCR clonality analysis demonstrated that Ag-Neo2/15 scaffolds promoted the expansion of functionally superior T-cell clones. The top clones of CD8^+^ T cells expanded with Ag-Neo2/15 scaffolds exhibited a favorable phenotype, essential for effective antigen recognition and sustained T-cell mediated cytotoxicity.

Our findings suggest that Ag-Neo2/15 scaffolds represent an advancement in ACT by producing high-quality, functional antigen-specific T cells. This method has the potential to improve clinical outcomes in cancer therapy by generating large numbers of highly functional T cells, thereby optimizing the balance between cytotoxicity and proliferation capacity with less exhausted T-cells in expansion protocols.

## Introduction

Adoptive cell therapy (ACT) relies on the infusion of *ex vivo* expanded T cells into cancer patients to generate an T cell-mediated antitumor response. ACT strategies can be broadly categorized into two approaches: I) T cells obtained from the patient’s blood, genetically engineered to express a tumor-specific T cell receptor (TCR) or a chimeric antigen receptor (CAR), and II) naturally occurring tumor-infiltrating lymphocytes (TILs) from surgically removed tumor samples. Engineered T-cell therapies have shown remarkable efficacy against hematological cancers (1, 2). Recently, engineered T-cell therapies have also shown efficacy in solid cancers (3, 4); however, the clinical benefit is often not sustained long-term due to tumor antigen heterogeneity or loss of targeted antigen (5). Additionally, in the setting of solid tumors, engineered T-cell therapies have caused severe and fatal toxicities due to cross-recognition of life-sustaining healthy tissue (5, 6). On the contrary, TILs have the potential to overcome tumor antigen heterogeneity by offering a polyclonal and multi-targeted approach. Additionally, TIL therapy is considered safer as compared to engineered T-cell therapies due to their maturation in the host, including a thorough thymic-selection process, limiting the risk of cross-recognition. TIL therapy has shown great promise as treatment for metastatic melanoma (7) leading to the recent FDA approval of Lifileucel (Amtagvi) for patients with advanced melanoma (8). Despite the advancements in TIL therapy, several limitations related to manufacturing needs to be addressed to harvest their full potential. Manufacturing of TILs relies on lengthy and non-specific *ex vivo* expansion, promoting terminal differentiation and T-cell exhaustion (9, 10). To address these challenges, we recently reported on the development of artificial antigen-presenting scaffolds (Ag-scaffolds) for specific *ex vivo* expansion of circulating tumor-specific T cells from peripheral blood (11). Ag-scaffolds are composed of a dextran-polysaccharide backbone containing multiple streptavidin molecules, enabling site-specific conjugation of biotinylated proteins such as peptide-MHC (pMHC), cytokines, and co-stimulatory molecules. Using Ag-scaffolds decorated with pMHCs, IL-2, and IL-21, we were able to generate polyclonal T-cell products enriched for antigen-specific T cells with high self-renewal capacity, low exhaustion, and enhanced cytotoxicity (11).

Recently, *de novo*-designed Neoleukin-2/15 (Neo2/15) emerged as a hyper stable protein mimicking the functional properties of IL-2 and IL-15 (12). Neo2/15 specifically binds to the shared IL-2Rβ/γ receptor subunits (CD122/CD132), involved in T-cell activation while excluding the IL-2Rα receptor (CD25), thus reducing the activation of regulatory T cells (Tregs). In pre-clinical models, Neo2/15 treatment induced strong proliferation of CD8^+^ T cells with limited expansion of Tregs, leading to overall improvements in anti-tumor efficacy and survival (12). Thus, this study investigates the expansion kinetics, molecular and cellular properties, and functional characteristics of T cells expanded for ACT using Ag-scaffolds loaded with pMHC and hyper-stable Neo2/15.

## Results

### Antigen-presenting scaffolds loaded with Neo2/15 expand functional T cells

To ensure site-specific conjugation of Neo2/15 to the dextran backbone, we designed a recombinant version of Neo2/15 carrying a C-terminal avitag (Neo2/15-avi). To test the functional activity of Neo2/15-avi, we used CFSE dilution to determine CD8^+^ T-cell proliferation. The activity of Neo2/15-avi was compared to recombinant IL-2. As the potency of Neo2/15-avi was unknown, we used molar concentration of Neo2/15-avi corresponding to 10, 100, and 1000 units of IL-2 for stimulation. Non-stimulated T cells were used as baseline, and T cells stimulated with anti-CD3/anti-CD28-coated Dynabeads and 20U/mL IL-2 as positive control. Eight days after stimulation, CFSE dilution was assessed by flow cytometry. Stimulation with 1000 U/mL of Neo2/15-avi or IL-2, respectively, resulted in similar levels of proliferation with more than 90% of CD8^+^ T cells having divided more than once (Figure 1A). Lower concentrations of IL-2 and Neo2/15-avi also resulted in a similar number of cell divisions (Supplemental Figure 1). Based on this, we concluded that Neo2/15-avi was fully functional and could be further assessed for expansion of antigen-specific T cells using Ag-scaffold technology. Ag-scaffolds are designed to ensure pMHC-directed expansion of antigen-specific T cells, while avoiding unspecific expansion of bystander T cells. To find the optimal molar ratio between dextran, pMHC, and Neo2/15-avi (dextran:pMHC:Neo2/15-avi), we assembled Ag-scaffolds with a constant pMHC amount, determined by the HLA type, and varying amounts of Neo2/15-avi relative to dextran. The resulting Ag-Neo2/15 scaffolds, had increasing molar ratio of Neo2/15 bound to the Ag-scaffold, here denoted Ag-Neo2/15 (Neo-1x), Ag-Neo2/15 (Neo-3x), Ag-Neo2/15 (Neo-6x), and Ag-Neo2/15 (Neo-12x). These scaffolds were supplemented to cell cultures every three to four days during a 14-day expansion period. Expansion with Ag-Neo2/15 scaffolds with higher Neo2/15-avi ratio resulted in significantly higher total numbers of cells after 14 days of culture (Figure 1C). On day 14 of expansion, we performed tetramer staining to determine the percentage of antigen-specific T cells in the cultures (Figure 1D). Although all cultures expanded with Ag-Neo2/15 scaffolds showed a significant increase in the percentage of antigen-specific T cells compared to baseline (∼1% antigen-specific T cells), we found the highest frequency of antigen-specific T cells in the Ag-Neo2/15 (Neo-3x) scaffold condition. However, as the amount of Neo2/15-avi attached to the Ag-scaffolds increased, the frequency of antigen-specific T cells progressively declined. When determining the total yield of antigen-specific T cells in the cultures we found the highest number of cells in cultures expanded with the Ag-Neo2/15 (Neo-3x) scaffold (Figure 1E). Next, we addressed the functionality of the expanded cells using flow cytometric analysis of intracellular cytokine staining. We observed a tight correlation between the overall frequency of triple-positive T cells (TNFα+/IFNγ+/CD107a+) and the percentage of antigen-specific T cells after co-culture with antigen-positive target cells (Figure 1G and 1F). We observed lower level of PD-1 expression on CD8^+^ T cells expanded with higher number of loaded Neo2/15 molecules on the scaffold, while all the expanded T cells consistently expressed TIM3 and LAG3 across scaffold condition (Figure 1H).

**Figure 1:**
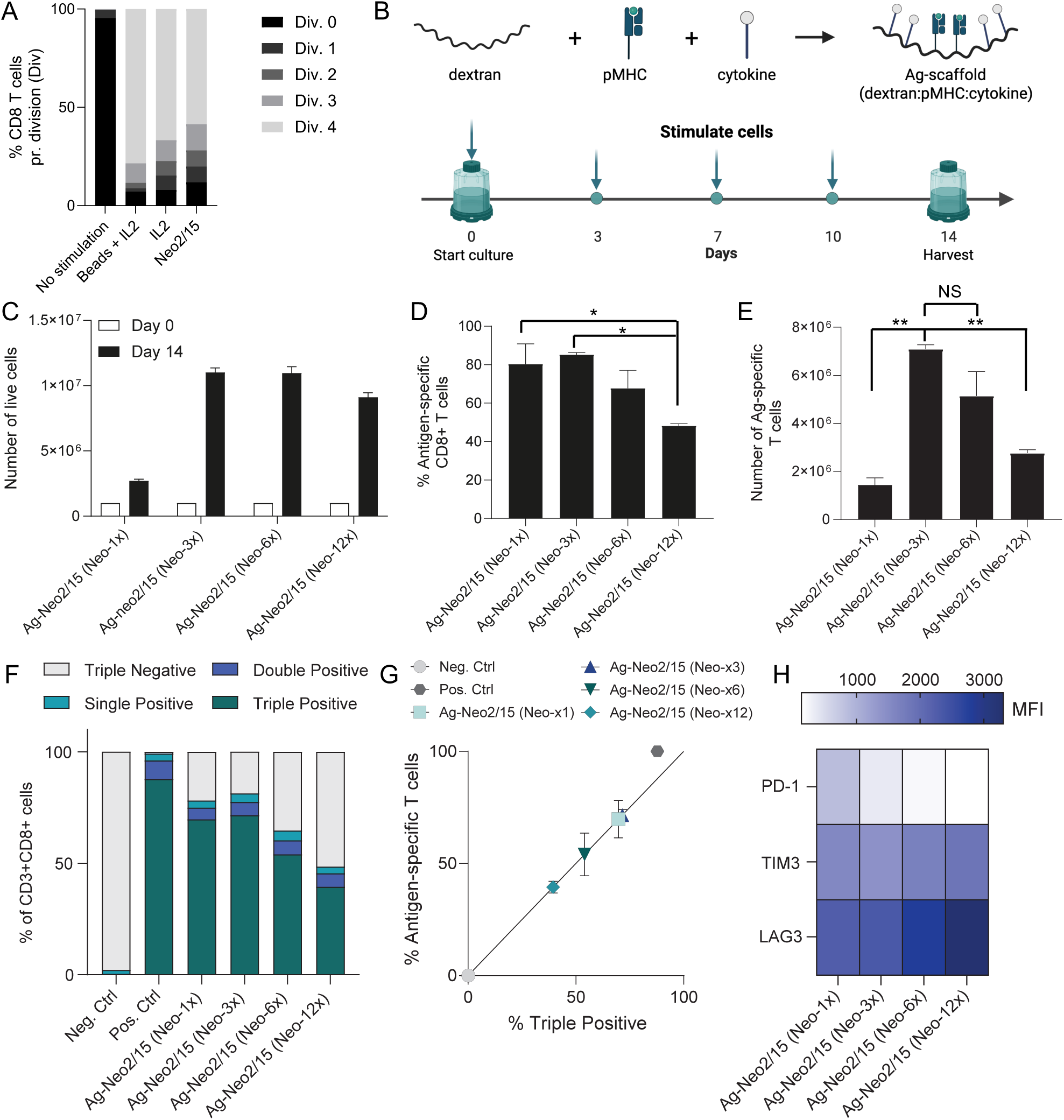
Neo2/15 induces strong proliferation and expansion of antigen-specific T cells. A) Recombinant human biotinylated Neo2/15 induces stronger proliferation of CFSE labeled naïve T cells compared to recombinant human IL-2. B) Schematic representation of artificial antigen-presenting Ag-scaffolds (top panel) and the experimental expansion setup (bottom panel). C) Number of live cells days after expansion with Ag-Neo2/15 scaffolds. D) Percent of antigen-specific T cells and E) number of antigen-specific (Ag-specific) T cells in culture after 10 days of expansion. F) The percentage of cells in culture expressing one, two, or three cytokines (CD107a, TNFa, IFNg) as determined by intracellular cytokine staining (ICS). G) The relationship between % Triple positive cytokine T cells and the % of antigen-specific cells in culture. H) The mean fluorescence intensity of exhaustion markers PD1, TIM3, and LAG3 on T cells expanded with Ag-Neo2/15 scaffolds determined by flow cytometry. Figure 1A was created in BioRender. Hadrup, S. (2025) https://BioRender.com/9kaiiwf.

Based on these findings, we concluded that Ag-Neo2/15 (Neo-3x) scaffolds were superior for expanding the highest level of antigen-specific T cells while maintaining a high functional profile dominated by triple cytokine-secreting T cells.

Next, we investigated the functional capacity of expanded T cells, in terms of killing target cells and production of the effector cytokine IFNγ. In this setup, we compared Ag-Neo2/15 scaffolds to IL-2/IL-21-loaded Ag-scaffolds with dextran:pMHC:IL-2:IL-21 ratio of 1:pMHC:6:6 (Ag-IL2/21 scaffolds) described by Tvingsholm *et al.* (10). We here selected Ag-Neo2/15 (Neo-3x) scaffolds because it demonstrated the best T cell expansion while maintaining a favorable functional profile, and Ag-Neo2/15 (Neo-12x) scaffolds to more closely resemble the cytokine quantity on the Ag-IL2/21 scaffolds. First, we measured the IFNγ release from expanded T cells after 24h of co-incubation with target cells. T cells expanded with Ag-Neo2/15 scaffolds produced more IFNγ as compared to T cells expanded with Ag-IL2/21 scaffolds (Figure 2A).

**Figure 2:**
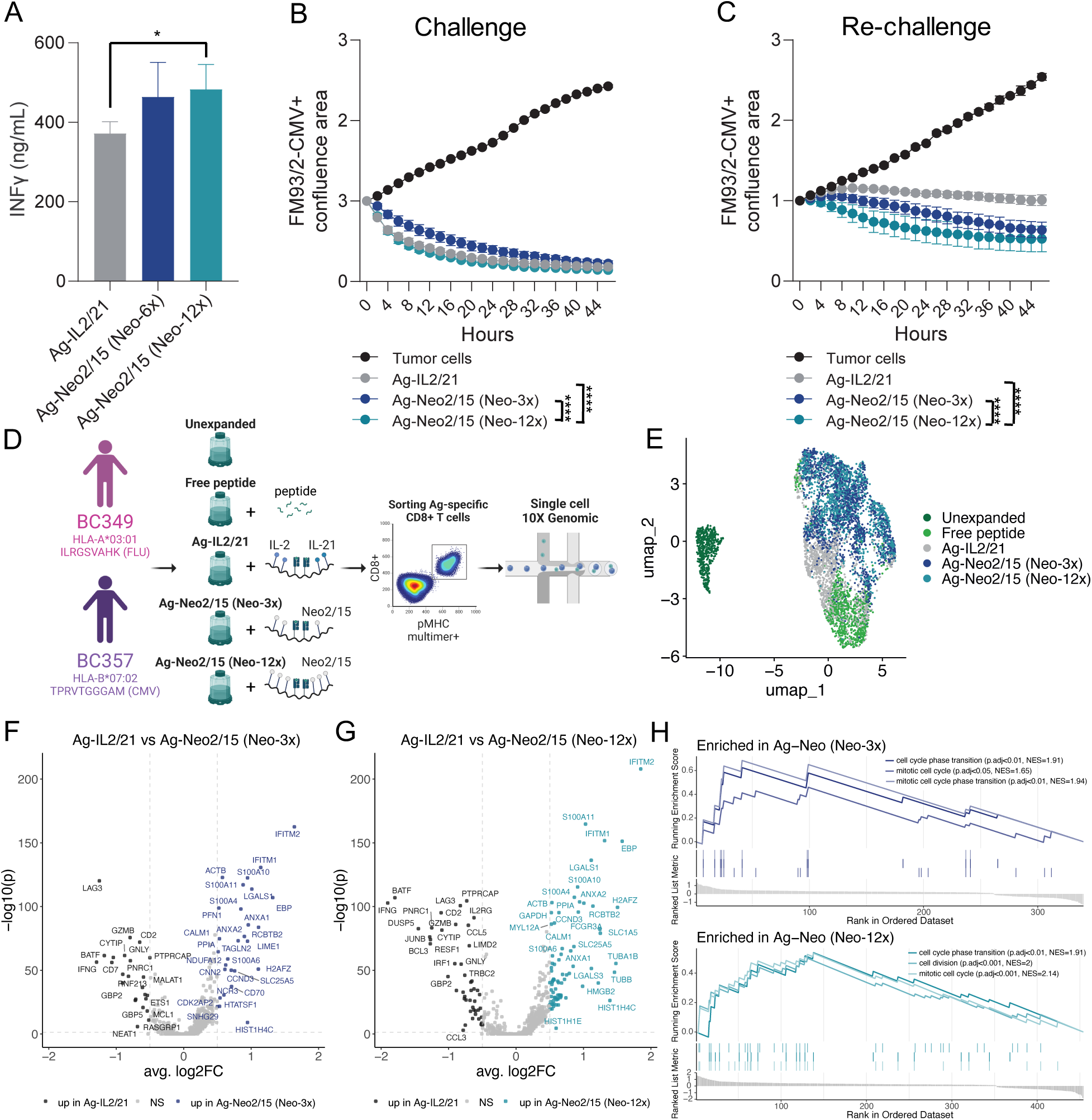
Antigen-specific CD8^+^ T cell expanded with Neo-2/15 exhibit a distinct transcriptomic profile. A) The level of IFNg measured by ELISA in cell culture supernatant of Ag-scaffold expanded T cells co-cultivated with target cells. B) The cytotoxicity of Ag-scaffold expanded T cells over a 44h co-culture period with target cells measured by Incucyte. C) The cytotoxicity of Ag-scaffold expanded T cells over a 44h co-culture period when re-challenged with target cells measured by Incucyte. D) Experimental workflow for the single-cell analysis of CD8+ and multimer+ T cells from human PBMCs under different *in vitro* expansion strategies. Using T cells from HLA-A*03:01 Influenza NP IRL-specific and HLA-B*07:02 CMV pp65 TPR-specific T cells from two healthy donors. E) UMAP projection of the scRNA-seq data colored based on the five different expansion strategies. F) Volcano plot showing key differentially expressed genes between the Ag-IL2/21 and Ag-Neo2/15 (Neo-3x) and G) between the Ag-IL2/21 and Ag-Neo2/15 (Neo-12x). H) Selected biological pathways determined by the gene set enrichment analysis when comparing Ag-IL2/21 and Ag-Neo2/15 (Neo-3x) (top panel) and when comparing Ag-IL2/21 and Ag-Neo2/15 (Neo-12x) (bottom panel). Figure 2B was created in BioRender. Hadrup, S. (2025) https://BioRender.com/9kaiiwf.

The increased production of IFNγ was accompanied by superior cytotoxic functionality of T cells expanded with Ag-Neo2/15 scaffolds, enabling them to effectively kill target cells (Figure 2B-C, Supplemental Figure 3). Notably, this killing capacity was even more pronounced during the rechallenge compared to the initial 46 hours of co-culturing with target cells (Figure 2C, Supplemental Figure 3).

### Single-cell analysis of antigen-specific T cells expanded with Ag-Neo2/15 scaffolds

After observing the enhanced killing capacity of CD8+ T cells expanded with Ag-Neo2/15 scaffolds, we performed a single-cell analysis using antigen-specific T cells from two healthy donors to further investigate differences between cells expanded with Ag-Neo2/15 scaffolds and those expanded using conventional strategies. In this analysis, we examined the transcriptome, surface proteome, and T-cell clonality of unexpanded T cells, as well as T cells expanded using free peptide, IL-2 and IL-21, Ag-IL2/21 scaffolds, and two different Ag-Neo2/15 scaffolds (Neo-3x and Neo-12x). The donor-specific T cells were cultured for two weeks and stimulated twice weekly until harvest (Supplemental Figure 2). At harvest, CD8+ pMHC multimer+ T cells were sorted based on donor-specific viral responses to either the Influenza NP ILR-peptide bound to HLA-A*03:01 or the CMV pp65 TPR-peptide bound to HLA-B*07:02 and analyzed by single-cell transcriptomics (Figure 2D).

UMAP analysis of the transcriptome revealed distinct clusters of T cells under the various expansion conditions (Figure 2E). Unstimulated cells formed a separate cluster (on the left), suggesting a unique transcriptional profile, while cells expanded with free peptide and cytokines, Ag-IL2/21 scaffolds, and the two Ag-Neo2/15 scaffolds (Neo-3x and Neo-12x) clustered closely, indicating less transcriptional differences. The same trend was observed when evaluating the differential expressed genes within each expansion condition (Supplemental Figure 4).

We have previously demonstrated the superiority of the Ag-IL2/21 scaffold when compared to more traditional expansion methods like free peptides and cytokines (10); therefore, we chose the Ag-IL2/21 scaffold as our most advanced expansion method for comparison to Ag-scaffolds loaded with Neo2/15, to investigate how the different cytokines affect the final T-cell products.

To investigate the difference between T cells expanded with Ag-IL2/21 scaffolds and cells expanded with Ag-Neo2/15 scaffolds, we performed a differential expression analysis (DEA) (Figure 2F and G), followed by a gene set enrichment analysis (GSEA) (Figure 2H, Supplemental Figure 5 and Supplemental Figure 6). T cells expanded with Ag-Neo2/15 scaffolds showed higher expression of genes linked to cell division and mitotic cell cycle (TUBA1B, TUBB, CCND3, CDK2AP2), indicating that these are more proliferative than T cells expanded with Ag-IL2/21 scaffolds. Interestingly, this also resulted in a higher number of antigen-specific CD8^+^ T cells in cultures expanded with Ag-Neo2/15 scaffolds compared to Ag-IL2/21 scaffolds (Supplemental Figure 2C). Meanwhile, T cells expanded with Ag-IL2/21 have a higher expression of genes linked to T cell effector functionality (CCL5, IL2RG, IFNG, IL2RG, BATF, IRF1).

Similar trends are evident when examining surface markers derived from the single cell analyses (Supplemental Figure 7 and Supplemental Figure 8). Notably, CD8^+^ T cells expanded with Ag-Neo2/15 scaffolds displayed higher expression of the inducible costimulatory receptor ICOS compared to those expanded with Ag-IL2/21 scaffolds. ICOS is expressed on activated T cells and plays a role in T cell proliferation, cytokine production, and survival. However, CD8^+^ T cells expanded with Ag-IL2/21 scaffolds exhibited elevated levels of markers associated with activation (CD69, CD7, CD2 and CD27) and exhaustion (CD39) (Supplemental Figure 7 and Supplemental Figure 8).

Interestingly, the expression of homing molecules differed between cells expanded with either Ag-IL2/21 or Ag-Neo2/15 scaffolds. CD8^+^ T cells expanded with Ag-Neo2/15 scaffolds showed higher expression of the intestinal homing molecule, alpha4-beta7 integrins (Integrin beta7 and CD49d); whereas those expanded with Ag-IL2/21 exhibited increased expression of more generic homing molecules, including CD103 (alphaE-beta7 integrin homing to epithelial tissues) and CD11a (forming LFA-1 together with CD18). This suggests that the different scaffolds can affect the expression of homing molecules on the expanded T cells, guiding them to specific tissue types.

Overall, our single-cell analysis revealed that T cells expanded with Ag-Neo2/15 scaffolds exhibited distinct transcriptional profiles and higher proliferation rates compared to those expanded with Ag-IL2/21 scaffolds. Ag-Neo2/15-expanded T cells showed increased expression of genes linked to cell division and higher levels of the inducible costimulatory receptor ICOS. In contrast, Ag-IL2/21-expanded T cells had higher expression of activation and differentiation genes, along with elevated levels of activation and exhaustion surface markers.

### T cells expanded with Ag-Neo2/15 scaffolds have superior functional characteristics

To assess the functional state of the different expansion conditions, we used the top 30 cytotoxicity (cyt) and dysfunctionality (dys) gene signatures, described by Li *et al.* (13), to calculate a combined score for each gene set. The resulting scores for each cell were visualized using UMAPs (Figure 3A), and the differences between these scores for each expansion strategy were quantified (Figure 3B).

**Figure 3:**
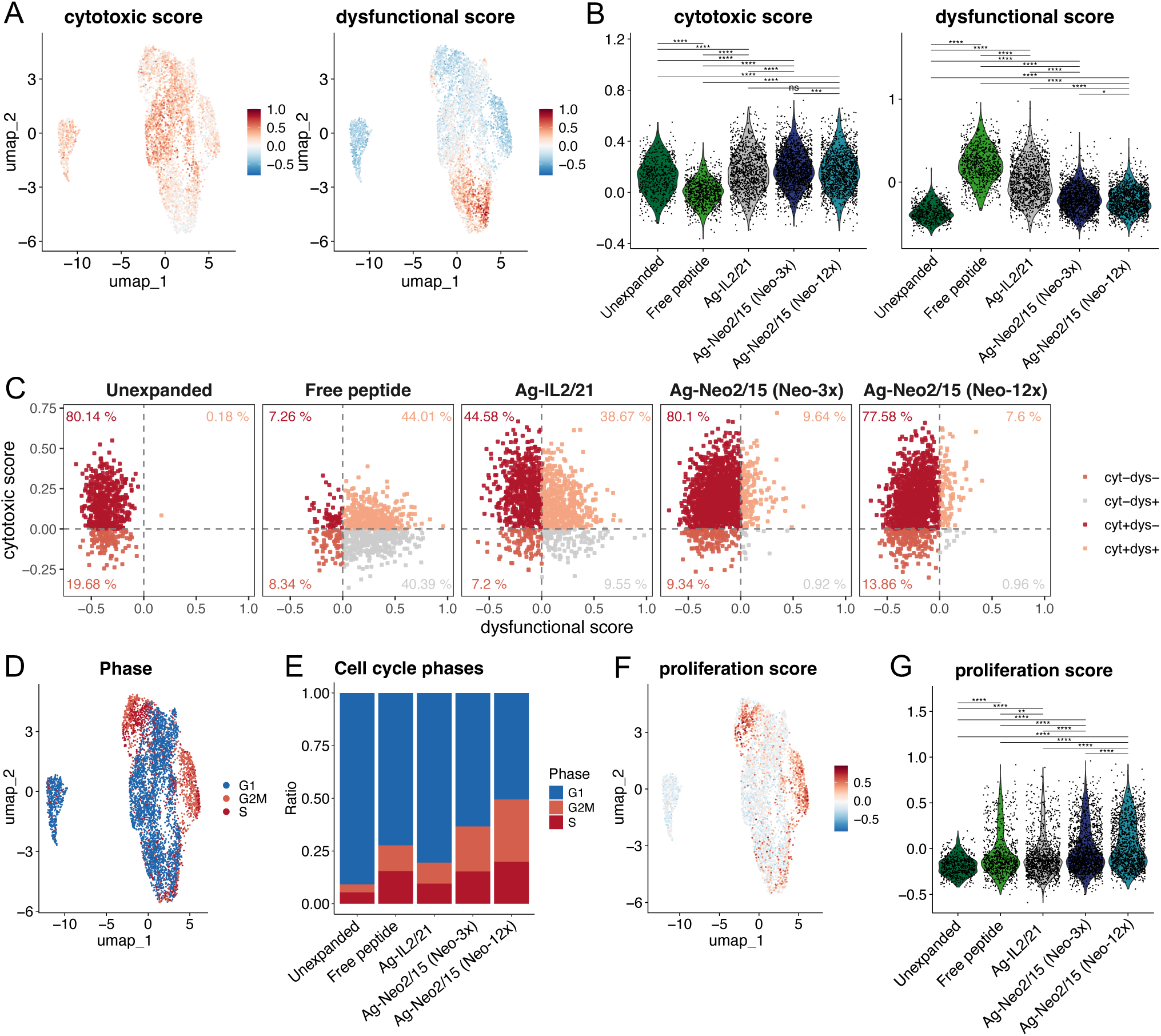
T cells expanded with NeoIL2/15 have a more favorable phenotype and shows tumor killing. A) UMAP shows the cytotoxic and dysfunctional score for each cell. B) Violin plots showing the expression level of the cytotoxic and dysfunctional score for each cell within each of the five expansion strategies. C) Quadrant scatter plots showing the relationship between the cytotoxic and dysfunctional score for each cell for each expansion strategy. Each scatter plot is divided into four quadrants. The four quadrants represent: cyt^+^dys^+^ (both scores > 0), cyt^-^ dys^-^ (both scores < 0), cyt^+^dys^-^ (cytotoxic > 0, dysfunctional < 0), and cyt ^-^ dys^+^ (cytotoxic < 0, dysfunctional > 0). D) UMAP showing cell cycle phase G1, G2M and S for each cell. E) Ratio of the three cell cycle phases for each expansion strategy. F) UMAP showing proliferation score for each cell. G) Violin plots showing the expression level of the proliferation score for each expansion strategy. The statistical test shown between the five different expansion strategies were compared using an unpaired, non-parametric Wilcoxon test of the mean expression, where p values > 0.05 were considered statistically non significant and represented as ns, whereas p ≤ 0.05, p < 0.01, p < 0.001 and p < 0.0001 is represented as *, **, *** and ****, respectively.

T cells expanded with Ag-Neo2/15 scaffolds showed significantly higher cytotoxicity scores compared to both unstimulated and free peptide-expanded cells, whereas T cells expanded with Ag-IL2/21 scaffolds had more comparable cytotoxicity scores. This suggests that cells expanded with Ag-Neo2/15 scaffolds induce a potent cytotoxic T-cell response similar to that seen with Ag-IL2/21 scaffolds (Figure 3B).

Furthermore, Ag-Neo2/15-expanded T cells exhibited significantly lower dysfunctionality scores compared to those expanded with either free peptides or Ag-IL2/21 scaffolds (Figure 3B). As expected, unexpanded cells had the lowest dysfunctionality scores, as these cells have not been expanded and, therefore, have not undergone repeated activation or stimulation that could lead to exhaustion or dysfunction.

To further investigate the relationship between the cytotoxicity and dysfunctionality score for each cell, we generated a scatter plot for each of the different expansion conditions (Figure 3C). This plot underscores the effect that the different expansion conditions have on the T-cell phenotype. Notably, Ag-Neo2/15-expanded cells have a higher proportion of both cyt^+^dys^-^ phenotypes, with fewer cells in the cyt^-^dys^+^ quadrants. This indicates that the Ag-Neo2/15 scaffold expansion not only promotes enhanced cytotoxicity but also preserves T-cell functionality by reducing the expression of dysfunction-associated genes.

These results highlight that expansion with Ag-Neo2/15 scaffolds maintain a cytotoxic phenotype while having lower dysfunctionality scores, distinguishing it from other expansion conditions. The higher cytotoxicity scores observed in Ag-Neo2/15-expanded T cells, suggest that this Ag-scaffold may provide a powerful tool for enhancing T-cell-based immunotherapies. Additionally, the reduced dysfunctionality scores in Ag-Neo2/15-expanded T cells indicate that this approach helps maintain T-cell functionality, potentially improving the efficacy of adoptive T-cell therapies. Based on the findings in Figure 1, which suggest that Ag-Neo2/15 scaffold expansion promotes a more proliferative state in T cells, we aimed to validate this observation using the single cell data. Looking into the cell cycle phase dynamics, we observe that most cells across all conditions remain in the G1 phase, where cells grow and prepare for DNA replication (Figure 3D and E). Smaller populations were observed in the S phase, where DNA replication occurs, and in the G2M phase, associated with cell division. However, the cell cycle phase distribution differed among the different expansion conditions (Figure 3E). Notably, Ag-Neo2/15-expanded cells had an increased proportion of cells in the S and G2M phases, suggesting that these cells are more proliferative. Furthermore, proliferation scores, calculated from the mean expression for a defined proliferation gene set, further confirmed this trend (Figure 3F and G). As seen on the UMAP and quantified in violin plots, cells expanded with Ag-Neo2/15 scaffolds consistently exhibit higher proliferation scores than those left unexpanded or expanded with free peptides or Ag-IL2/21 scaffolds.

Collectively, these results suggest that Ag-Neo2/15 scaffold expansion promotes a highly proliferative, cytotoxic T cell with reduced dysfunctionality and superior functional capacity, supporting its potential application in adoptive cell therapies.

### Ag-Neo2/15 scaffolds expand T cell clones with superior functional characteristics

To further elucidate differences between the T-cell products generated under the various expansion conditions, we analyzed the TCR clone distribution for each condition (Figure 4A). This analysis revealed that the expansion of TCR clones was largely similar across expansion conditions (Figure 4A). However, some variation was observed especially for BC357 indicating that the clonal expansion can be affected by the cytokine loaded onto the scaffolds. To investigate the similarity of these clones, we compared the CDR3 alpha and beta sequences for the top four T-cell clones per donor and showed that these sequences are quite distinct and differs in both length and minimal sequence similarity (Figure 4B, Supplemental Table 1). Even though the T cells from each donor only recognize a single peptide-MHC complex, these T cells maintain a diverse TCR repertoire, an observation that aligns with previous findings (14). Looking further into the phenotypic characteristics of the four most dominant TCR clones for each donor, we found that the top clones expanded with Ag-Neo2/15 scaffold had a higher proportion of cyt^+^dys^-^ (Figure 4C, Supplemental Figure 9). This indicates that both the phenotype and clonality of the clones are influenced by the different expansion conditions.

**Figure 4:**
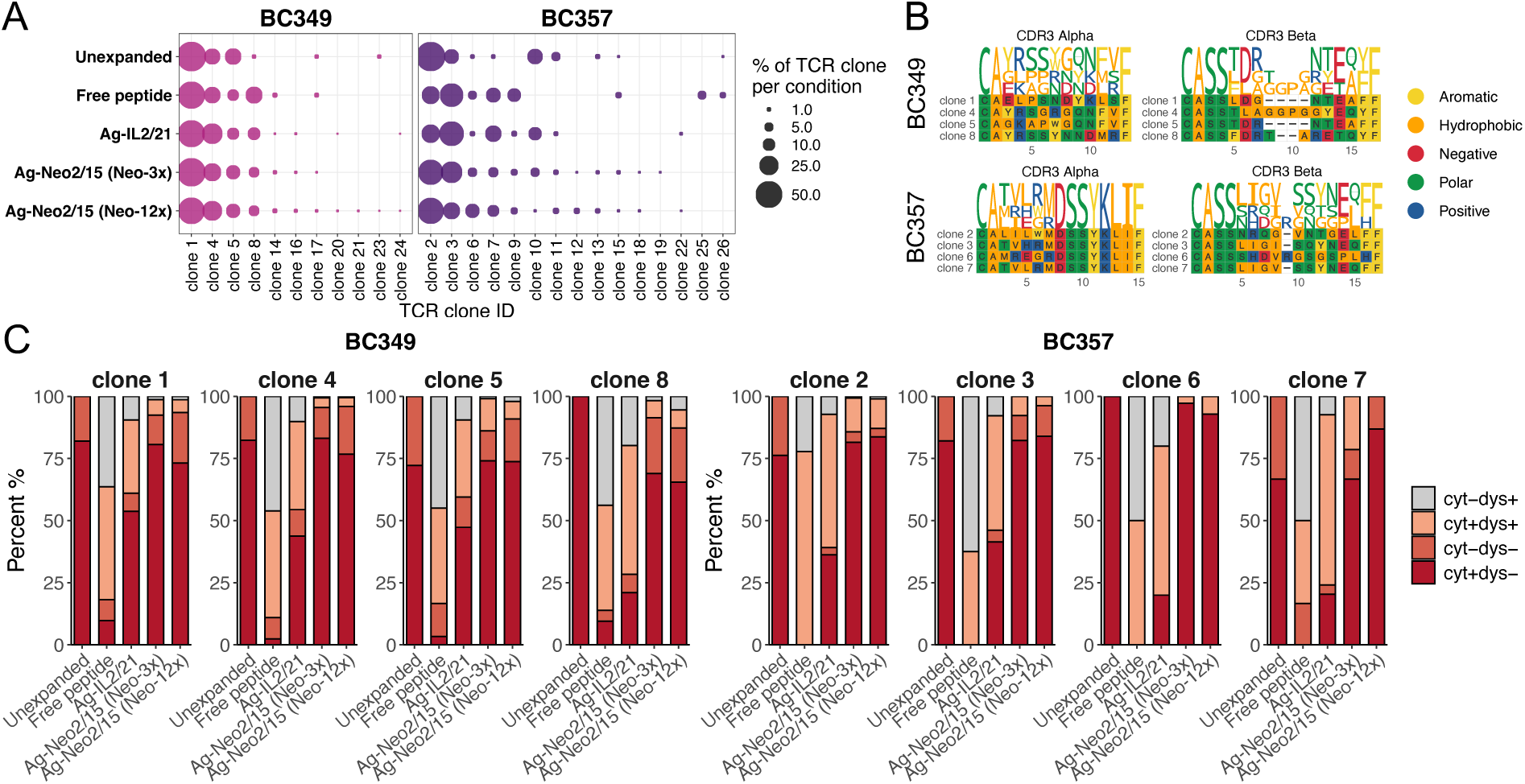
TCR diversity and clonal expansion. A) TCR clonotype distribution of the antigen-specific T cells from the single cell experiment. Circle size corresponds to clonal frequency out of all TCR clones detected within each expansion strategy. B) UMAPs of the top four TCR clonotypes for each of the two healthy donors, BC349 (left) and BC357 (right). C) Distribution of the relationship between the cytotoxic and dysfunctional score for each of the five expansion strategies within the top four TCR clonotypes for each of the two donors. The four colors represent: cyt^+^dys^+^ (both scores > 0), cyt ^-^dys^-^ (both scores < 0), cyt^+^dys^-^ (cytotoxic > 0, dysfunctional < 0), and cyt ^-^dys^+^ (cytotoxic < 0, dysfunctional > 0).

## Discussion

This study demonstrates the successful development and optimization of an antigen-specific T cell expansion platform utilizing Neo2/15, an engineered neoleukin IL-2/IL-15 chimera, conjugated to dextran-based antigen scaffolds. Our findings reveal significant advantages of this system for generating high-quality antigen-specific T cells for adoptive cell therapy applications. The results show that Neo2/15 has the ability to stimulate CD8^+^ T-cell proliferation comparable to recombinant IL-2 across various concentrations. This finding is noteworthy as Neo2/15 is engineered to reduce side effects commonly associated with IL-2 therapy, such as vascular leak syndrome (VLS), while maintaining potent immunostimulatory properties (15). The comparable proliferation rates observed in our study confirm that Neo2/15 can be a relevant alternative to IL-2 in T-cell expansion protocols. Building on this validation, we incorporated the recombinant Neo2/15 to previously established antigen scaffolds to expand antigen-specific CD8^+^ T cells. Importantly, CD8^+^ T cells expanded with the optimized Ag-Neo2/15 scaffolds maintained high functionality, as assessed by cytokine production (TNFα, IFNγ) and degranulation (CD107a) upon antigen re-stimulation.

Our findings demonstrate that T cells expanded with Ag-Neo2/15 scaffolds exhibit superior functional characteristics compared to those expanded with Ag-IL2/21. Notably, Ag-Neo2/15-expanded T cells produced significantly higher levels of IFNγ upon co-incubation with target cells, indicating an effector response. The killing assay results revealed that these T cells maintained superior cytotoxicity against target cells, even upon rechallenge. This enhanced killing capacity underscores the potential of Ag-Neo2/15 scaffolds to generate highly effective T cells for adoptive cell therapy, capable of sustained activity in immunosuppressive environments. Previous studies have demonstrated the potential of Neo2/15 in preclinical models, suggesting that Neo2/15 can significantly enhance the cytotoxic capacity of T cells, making them more effective in targeting and eliminating tumor cells (12, 16).

To further investigate T cells expanded using Ag-Neo2/15 scaffolds and compare them with other expansion strategies, including our recently established Ag-scaffold technology Ag-IL2/21 (11), we performed single-cell analysis on CD8^+^ T cells. This setup included unexpanded T cells and T cells expanded with either free peptide and cytokine (a conventional expansion method) or Ag-scaffolds decorated with pMHC, IL-2, and IL-21, as described previously by our group (11). Our transcriptomic analysis revealed differences between expanded CD8^+^ T cells using Ag-Neo2/15 scaffolds and those expanded by other methods. Notably, CD8^+^ T cells expanded with Ag-Neo2/15 scaffolds at both stoichiometries (Neo-3x and Neo-12x) exhibited considerable overlap in their transcriptional profiles, suggesting that the concentration difference had minimal impact on the overall transcriptional state of these cells. Differential expression analysis followed by GSEA indicated that CD8^+^ T cells expanded with Ag-Neo2/15 scaffolds showed increased expression of genes linked to cell division and proliferation capacity. Furthermore, cell cycle analysis demonstrated that Ag-Neo2/15-expanded cells had an increased proportion of cells in the S and G2M phases, suggesting enhanced proliferative capacity. This aligns with recent studies highlighting the importance of promoting T-cell proliferation while maintaining effector functions in adoptive cell therapy (17, 18). The ability to generate large numbers of highly functional T cells is a key factor in the success of cellular immunotherapies. This gene signature suggests that Ag-Neo2/15 scaffold expansion more effectively promotes T cell proliferation. The ability to induce robust proliferation while maintaining functional capacity is a crucial factor in developing effective T-cell based therapies.

Our immunophenotyping analysis on the single cell platform suggests that different expansion conditions significantly influence the homing characteristics and potential functionality of CD8^+^ T cells. Ag-Neo2/15 scaffold expanded cells indicate a preference for intestinal homing (α4β7 integrins), while Ag-IL2/21 scaffold expanded cells express more general homing molecules (CD103, CD11a/LFA-1). This distinction could have significant implications for the tissue tropism and migratory behavior of these expanded CD8^+^ T cells. The Ag-Neo2/15 scaffold expanded cells might be more suited for targeting intestinal tissues, whereas the Ag-IL2/21 expanded cells could have broader tissue-homing capabilities, including epithelial tissues. Moreover, the higher expression of ICOS on Ag-Neo2/15 scaffold expanded cells indicates a more costimulatory phenotype, which could lead to enhanced T-cell activation and function. ICOS is the hallmark of CD8+ tissue-resident memory T cells and is induced by T-cell activation and promotes T-cell proliferation and survival (19).

Our single-cell RNA sequencing analysis also revealed that T cells expanded with Ag-Neo2/15 scaffolds exhibited more favorable characteristics compared to the Ag-IL2/21 scaffold and free peptide expansion methods. Cells expanded with Ag-Neo2/15 scaffolds showed significantly higher cytotoxic scores and lower dysfunction scores, indicating a potent effector phenotype with reduced exhaustion markers. This aligns with our killing assay results, which demonstrated that T cells expanded with Ag-Neo2/15 scaffolds show enhanced functional killing capacity compared to those expanded with Ag-IL2/21 scaffolds. This finding is particularly important in the context of adoptive T-cell therapy, where maintaining T-cell functionality is crucial for therapeutic efficacy (17).

Our analysis of TCR clonality revealed that using the Ag-Neo2/15 scaffold expansion method could alter the distribution of dominant TCR clones. Moreover, the functional characteristics of expanded clones were markedly improved when expanded with Ag-Neo2/15 scaffolds, showing a higher proportion of cytotoxic and functional T cells. This suggests that our expansion method using Ag-Neo2/15 scaffolds promotes expansion of the T clones with enhanced functional quality, a crucial factor in maintaining diverse antigen recognition capabilities. The observation of distinct CDR3 sequences among the top clones, despite their identical antigen specificity, aligns with previous findings on TCR diversity in antigen-specific responses (14). This diversity likely contributes to a more robust immune response by allowing for varied recognition strategies of the same antigen, potentially enhancing the overall efficacy of the T cell product (17).

Our findings have significant implications for the field of adoptive T-cell therapy. The Ag-Neo2/15 scaffold expansion method offers a promising approach for generating large numbers of functional antigen-specific T cells. This approach has the potential to enhance the production of high-quality T cells for adoptive immunotherapy applications, potentially leading to improved clinical outcomes in cancer and infectious disease treatments.

## Methods

### Cell lines and healthy donor T cells

The HLA-A*0201-positive FM93/2 (ESTDAB-007) melanoma tumor cell lines were purchased from the European Searchable Tumor Cell Line Database (ESTDAB) and cultured as per supplier recommendation in RPMI media containing 10% FBS and 1% pen-strep. To establish a CMV CMV pp65 NLVPMVATV positive cell line, FM93/2 cells was lentivirally transduced with a bicistronic vector encoding click beatle green luciferase (CBG), mCherry fluorescent reporter, mouse mammary tumor virus (MMTV) gp70 signal peptide followed (20) by the NLVPMVATV peptide sequence (FM93/2-CMV+). FM93/2-CMV+ cells were single cells sorted to establish a clonal cell line. As a negative control, we generated an FM93/2 cell line solely expressing CBG-Luc and mCherry (FM93/2-CMV-). Peripheral blood mononuclear cells (PBMCs) were obtained from healthy donor buffy coats collected after informed consent and in accordance with the declaration of Helsinki from the central blood bank at Copenhagen university hospital (Rigshospitalet). Prior to expansion, HLA typing was performed, and T-cell reactivity was assessed against a panel of viral epitopes using DNA-barcoded pMHC multimers (11).

### Production of avitagged, biotinylated Neo2/15

The DNA sequence encoding Neo2/15 (12) was synthesized and cloned into the pET-30 plasmid between an N-terminal His-tag and C-terminal avitag for protein expression in *E.coli* (Neo2/15-avi) (Genscript). The E. coli strain BL21 (DE3) was co-transformed with plasmid and BirA for recombinant protein production. Briefly, a single colony was grown in TB media at 37 °C until an OD600 value of 1.2 was reached followed by isopropyl β-D-1-thiogalactopyranoside (IPTG) induction at 15°C for 16h. E. coli was pelleted by centrifugation and the supernatant collected for purification. Recombinant Neo2/15-avitagged protein was purified by one-step purification using a Ni-NTA column and sterilized by a 0.22μm filter before being stored in aliquots. The concentration was determined by BCATM protein assay with BSA as standard. The protein purity and molecular weight were determined by standard SDS PAGE along with Western blot confirmation. Neo2/15-avi was biotinylated using BirA biotin-protein ligase standard reaction kit (Avidity, LLC Aurora, Colorado) and purified with size exclusion chromatography using HPLC (Waters Corporation, USA). The protein concentration was measured using the Bradford assay, with BSA as the standard. Biotinylation efficiency was assessed via a gel-shift assay: briefly, biotinylated Neo2/15 forms multimers upon coupling with streptavidin, which were analyzed using standard SDS-PAGE.

### pMHC, tetramer and Ag-scaffold production

Relevant peptides were purchased from pepscan (Pepscan Presto BV), dissolved to 10 mM in DMSO and stored at −20°C until use. Recombinant Human Leukocyte Antigen (HLA) heavy chains and human β2 microglobulin light chain were produced in *E.coli*. We used a functionally empty disulfide-stabilized peptide-receptive HLA-A*02:01 variant (21), which can be loaded with peptide of interest during a 15-minute incubation at room temperature. For HLA-A*01:01 and HLA-B*07:02, the heavy and light chains were refolded with ultraviolet (UV)-sensitive ligands and purified as previously described (11) and pMHC complexes were generated by UV-mediated exchange with relevant peptides.

Tetramers were generated by conjugating pMHC monomers and fluorochrome labeled streptavidin. Blocking of free streptavidin sites was done with 25 uM D-Biotin (Avidity, BIO-200). Ag-scaffolds were assembled, using a streptavidin-conjugated dextran backbone (500 kDa) (Fina Biosolutions) mixed with biotinylated relevant pMHC molecules and Neo2/15-avi or avitagged IL-2 (Acro IL-2H82F3) and IL-21 (Acro IL-21-H82F7). Potential free streptavidin sites dextran backbone was blocked with D-biotin. To remove unbound molecules, Ag-scaffolds were filtered through 100kDA columns (Vivaspin6, Sartorius). Ag-scaffolds were stored at −80°C in a 10x freezing solution containing 5% Bovine Serum Albumin (BSA) and 50% Glycerol.

The dextran:pMHC ratio was experimentally determined for each HLA. The ratio for Ag-scaffold with HLA-A*02 was 1:36 or 1:24 depending on the batch, HLA-B*07 was 1:24 and HLA-A*03 was 1:24.

### Expansion of antigen-specific T cells using Ag-scaffolds

Specific T cells with pre-defined viral responses were expanded from PBMCs over a 10 to 14-day period. Briefly, PBMCs were cultured in VIVO 15 media (Lonza, BE02-060Q) supplemented with 5% human serum (Sigma Aldrich Heat Inactivated H3667) supplemented with Ag-scaffold (0.16 nM prefiltration) or specific peptide (5 µM), soluble IL-2 (40 U/mL, Peprotech 200–02) and soluble IL-21 (25 ng/mL, Peprotech 200–21). Ag-scaffold or free components were supplemented to the culture at initiation and twice weekly until harvest.

### T-cell cytotoxicity assay

The cytotoxic potential of Ag-scaffold expanded T cells was measured using the IncuCyte (Satorious). FM93/2-CMV+ cells or FM93/2-CMV-control cells were plated at a density of 5,000 cells/well and allowed to attach overnight. The following day, Ag-scaffold expanded T cells were added in an effector: target ratio of 1:1, and the plate was placed in the IncuCyte instrument. Data recording was set to 2-hour intervals over two days, and changes in impedance were reported as confluence areas. After the initial two-day measurement, the plate was removed from the Incucyte, cells were pelleted, and cell culture supernatant was collected for IFNγ ELISA. The pelleted cells were re-suspended in fresh medium and transferred to a new plate containing 5,000 FM93/2-CMV+ cells or FM93/2-CMV-control cells. Both FM93/2-CMV+ cells or FM93/2-CMV-control cells were plated 24h prior to rechallenge. The plate containing target and effector cells were placed in the IncuCyte and T-cell killing was followed for an additional two-day period.

### Intracellular cytokine staining

Expanded antigen-specific T cells were incubated with peptide-pulsed T2 target cells (E:T 1:1) for 4 hours in X-VIVO 15 media supplemented with αCD107a-PE antibody (BD 555801) and GolgiPlug (BD 550583). Co-incubation of antigen-specific T cells with non-pulsed T2 cells was used as negative control and co-incubation of antigen-specific T cells with non-pulsed T2 cells and leucocyte activating cocktail (LAC, BD, 550583) was used as positive control. Harvested Tcells were stained with LIVE/DEAD Fixable Near-IR cell marker (Invitrogen) for dead cell exclusion, CD3-FITC (BD 345764) and CD8-BV480 (BD 566121), fixed and permeabilized (Invitrogen 88882400) and stained with TNFα-PE-Cy7 (Biolegend 502930) and IFNγ-APC (BD 341117) and analyzed by flow cytometry.

### IFNγ ELISA

IFNγ in culture supernatants was monitored by ELISA. Briefly, wells were coated with (1µg/mL) anti-IFNγ capture antibody (BD 551221) prior to the addition of cell culture supernatant. The presence of IFNγ was detected using biotinylated 0,5µg/mL anti-IFNγ antibody (BD 554550). The reaction was visualized by a peroxidase-streptavidin conjugate (1:10,000; Merck 11089153001) and TMB (Kementec 4380A).

### 10xGenomics single-cell analysis platform

Cryopreserved PBMCs were thawed and stained with pooled DNA-barcoded pMHC multimers, TotalSeq™-C Human Universal Cocktail V1.0 (Biolegend), TotalSeq-C anti-human Hashtag antibodies for each distinct sample (BioLegend), LIVE/DEAD Fixable Near-IR cell marker (Invitrogen), and flow cytometry antibodies in presence of Human TruStain FcX Fc Blocking reagent (Biolegend) to block Fc receptors. The flow cytometry antibody mix was composed of CD8-BV480 (BD 566121, clone RPA-T8) and lineage antibodies: CD4-FITC (BD 345768), CD14-FITC (BD 345784), CD19-FITC (BD 345776), CD40-FITC (Serotech MCA1590F), CD16-FITC (BD 335035). The stained cells were next sorted on a FACS Melody (BD) to be run on 10xGenomics platform. We utilize the 10xGenomics 5′ v2 chemistry that allows us to use the immunophenotyping antibodies and DNA-barcoded pMHC multimers. Downstream processing of mRNA and DNA barcodes is performed according to the manufacturer’s instructions (Chromium Next GEM Single Cell 5’ Reagent Kits v2 [Dual Index], with the Feature Barcode technology for Cell Surface Protein & Immune Receptor Mapping) (10x Genomics, USA). The TCR, barcode (including pMHC multimers, immunophenotyping antibodies, and hashtag antibodies), and gene expression libraries were sequenced by Novogene on a NovaSeq running a 150 paired-end program.

### Single-cell data processing

The raw FASTQ files were processed using the CellRanger multi pipeline (10x Genomics, V.6.1.1), as this enables simultaneous profiling of the T-cell V(D)J repertoire, cell surface proteins (TotalSeq™-C), sample hashing, and gene expression. Both the V(D)J and gene expression were aligned to the GRCh37 reference. The filtered gene-barcode matrices from CellRanger were used for further analysis with Rstudio (version 4.3.2) and the Seurat package (version 5.0.2). Looking only at the sample hashing the data was first normalized using NormalizeData (normalization.method = “CLR”) and then demultiplex using HTODemux (positive.quantile = 0.99) and only cells classified as Singlet were used for the downstream analysis.

For the analysis of gene expression, genes identified in more than 3 cells and cells containing over 200 detected genes were included. To filter out low-quality cells, the following criteria were applied: 1) fewer than 6,000 detected genes per cell, 2) fewer than 25,000 molecules identified per cell, and 3) greater than 5% of Unique Molecular Identifiers (UMIs) originating from mitochondrial DNA. Following the removal of these low-quality cells, the SCTransform function was used for normalization of the gene expression matrices. Cell cycle effects were mitigated by regressing out S phase and G2/M phase scores during the data scaling process. Following this filtration, we had a total of 5,606 cells; 3,494 cells for donor BC349 and 2,112 cells for donor BC357 were included in further analyses. Additionally, donor-specific variations (BCs) were minimized through integration using the Harmony function from Seurat.

We previously reported the comparison between the unexpanded, free peptide, and Ag-IL2/21 scaffold expansion strategies (11). However, in this study, we have expanded our dataset by incorporating the two Ag-Neo2/15 scaffolds (Neo-3x and Neo-12x) into the single cell data. This addition allows for a comprehensive comparison of the Ag-Neo2/15 scaffolds with the previously used expansion strategies.

### Gene set enrichment analysis

The differential expressed genes, found between either Ag-IL2/21 and Ag-Neo2/15 (Neo-3x) or Ag-Neo2/15 (Neo-12x) scaffolds, were ranked by their average log2 fold changes and a gene set enrichment analysis (GSEA) was performed with the gseGO function from clusterProfiler (version 4.10.1) (18), using only biological process (ont=’BP’) with a minimum gene set above 10 (minGSSize = 10), and with a p-value cutoff below 0.05 (pvalueCutoff = 0.05). The resulting pathways were further filtered to have a qvalue < 0.1 and a GeneRatio > 0.5, and the enrichment score for each of these pathways are shown in Supplemental Figure 3. GSEA plots for selected enriched pathways were visualized using the gseaplot2 function from the enrichplot package in R.

### Gene enrichment scores

To evaluate the effect of multiple genes on a single cell level, we used the AddModuleScore function from Seurat to calculate enrichment scores for different gene sets, namely the cytotoxic, dysfunctional, and proliferation scores. The cytotoxic and dysfunctional score was calculated using the top30 cytotoxic and dysfunctional genes identified by Li *et. al.* (13). Here the dysfunctional gene set consists of LAG3, HAVCR2, PDCD1, PTMS, FAM3C, IFNG, AKAP5, CD7, PHLDA1, ENTPD1, SNAP47, TNS3, CXCL13, RDH10, DGKH, KIR2DL4, LYST, MIR155HG, RAB27A, CSF1, CTLA4, TNFRSF9, CD27, CCL3, ITGAE, PAG1, TNFRSF18, GALNT1, GBP2, and MYO7A, and the cytotoxic gene set includes IL7R, GNLY, FGFBP2, CX3CR1, FCGR3A, S1PR5, PLAC8, FGR, C1orf21, SPON2, CD300A, TGFBR3, PLEK, STK38, S1PR1, EFHD2, KLRF1, C1orf162, SORL1, FCRL6, TRDC, EMP3, CCND3, KLRG1, BIN2, SELL, KLRB1, SAMD3, and ARL4C.

The proliferation score was calculated using a gene set with the following genes, CCND3, CDK2, CDK4, CDK6, MKI67, E2F1, E2F2, PCNA, MCM2, MCM3, MCM4, MCM5, MCM6, MCM7, TOP2A, MYC, BCL2 and BCL2L1.

### T-cell receptor analysis

For the TCR analysis, we used only cells with at least one productive TCR α-chain and/or one productive TCR β-chain, selecting only the chain with the highest UMI count and read count if multiple chains were found within a cell. The clonotype ID was then manually annotated based on unique CDR3 sequences from the α- and β-chain. The CDR3 α and β sequences of the top four TCR clones were aligned using the imgt_align_junctions function from the cdr3tools library (https://github.com/caparks2/cdr3tools). The multiple sequence alignment and logo plot was performed using the ggmsa function from the ggmsa library (22).

### Phenotype analysis

The cell surface protein analysis was performed with cell surface proteins from the TotalSeq-C library, and the phenotype data were normalized and scaled using the NormalizeData and ScaleData function from Seurat.

### Differential expression analysis

To find differential expressed genes or phenotype makers between either Ag-IL2/21 and Ag-Neo2/15 (Neo-3x) or Ag-Neo2/15 (Neo-12x) scaffolds, the FindMarkers function from Seurat was used, with the logfc.threshold = 0 and min.pct = 0.75. Differential expressed genes or phenotype makers were shown using volcano plots generated with ggplot2 and heatmaps of all genes with an average log2 fold change above 0.5 and a p-value below 0.05 was generated with the ComplexHeatmap package.

## Authors contributions

MO, KKM, MK and SRH designed the research and wrote the manuscript. MO, KKM, MK, ST, KR, AKB, GK, GNA, GN, TT, SN, MSF and MK contributed to experiments and analysis. HRH, KHJ, MK and SRH provided important materials, methods and advice. SRH is responsible for the overall content as the guarantor. All authors read and approved the manuscript.

## Data availability statement

All data relevant to the study are included in the article or uploaded as supplementary information.

---

**Supplementary Figure 1:**
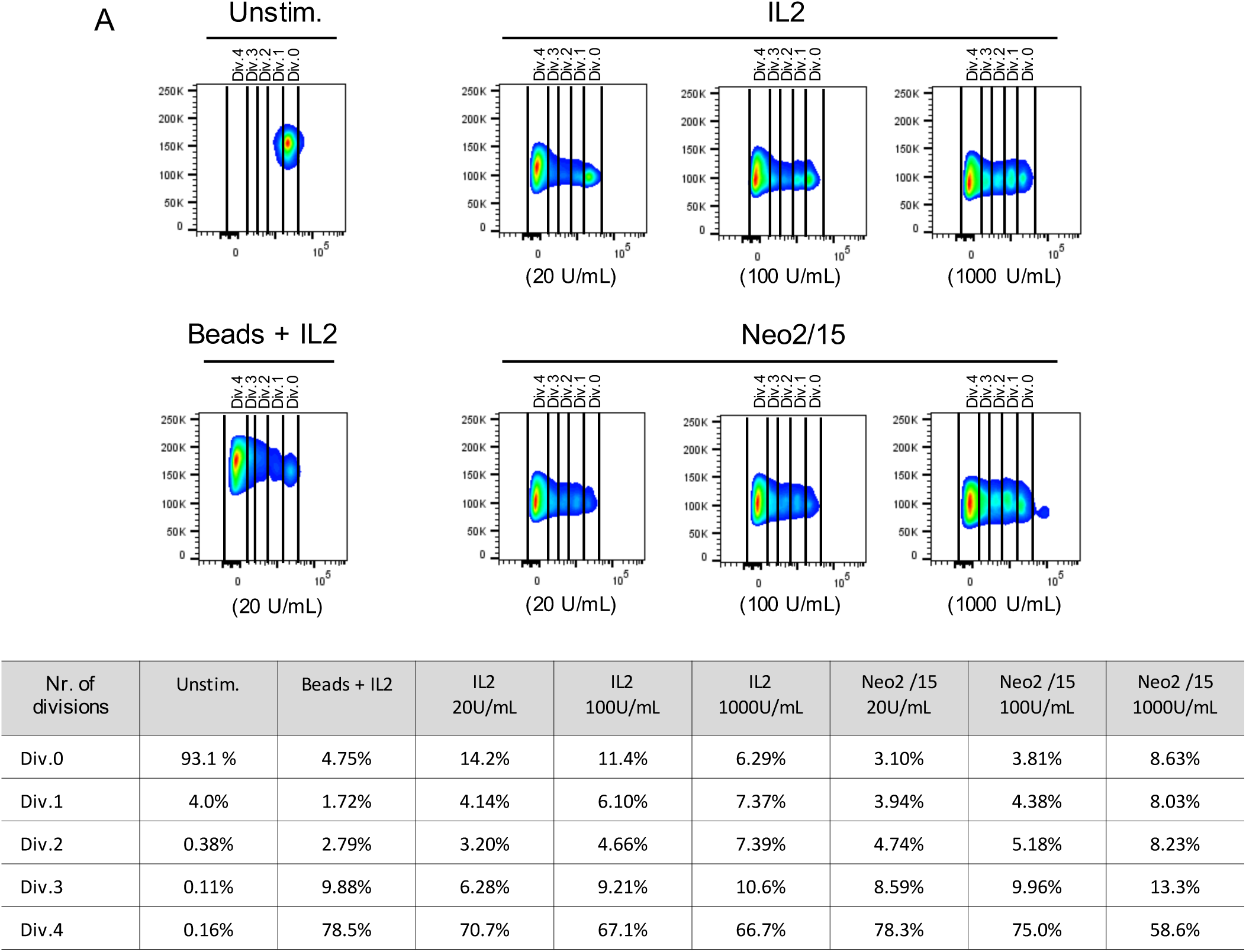
T cells stimulated with Neo2/15-avi and IL2 proliferate to a similar level. A) Flow plots showing the division of CD8+ T cells stimulated with molar ratios of IL2 or Neo2/15-avi as seen by dilution of CFSE cell trace dye. Unstimulated T cells were used a negative control and T cells stimulated with Dynabeads and Il2 as positive control.

**Supplementary Figure 2:**
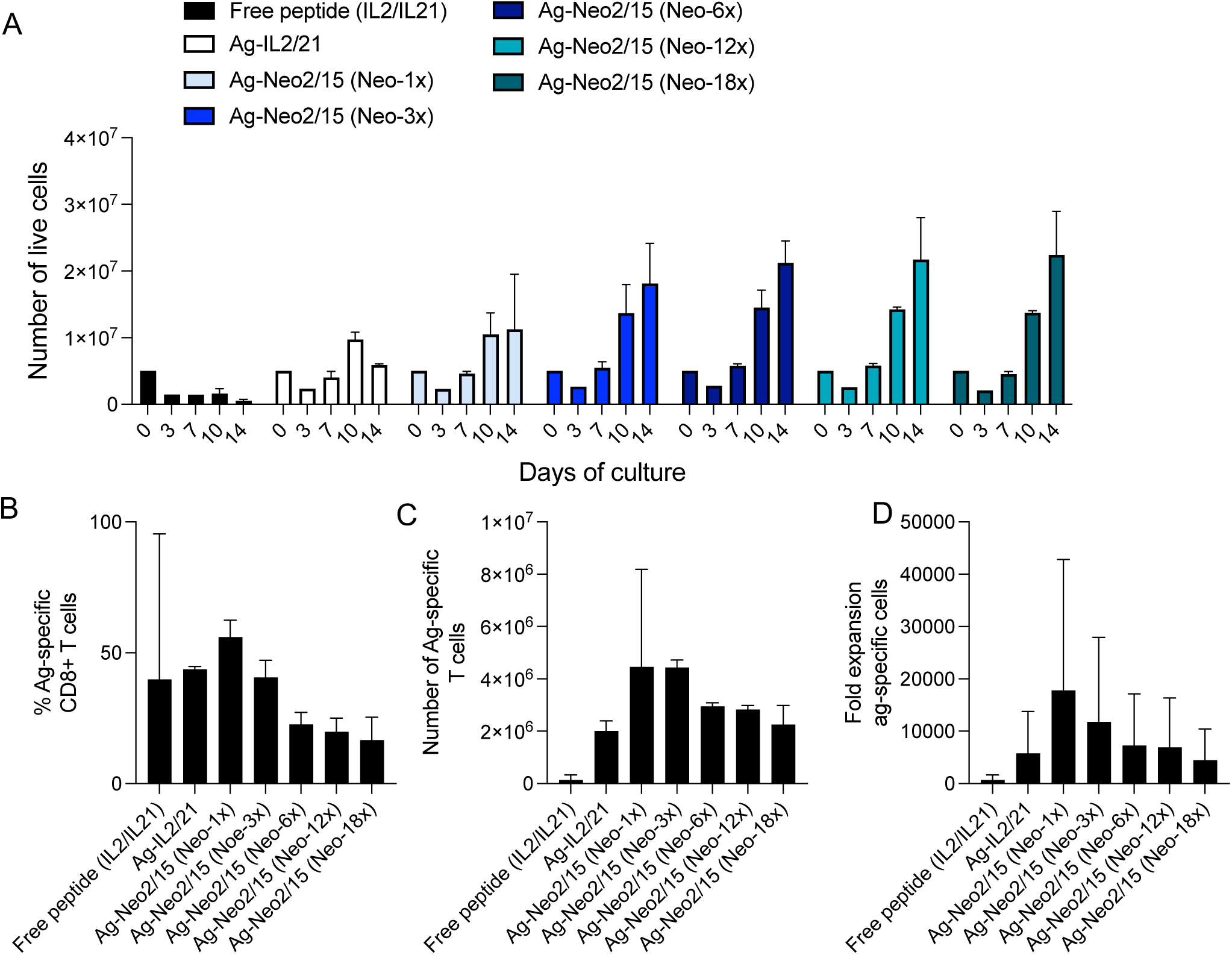
Expansion kinetics of T cells expanded with Ag-scaffold technology prior to single cell analysis. A) Number of live cells after a 14 day expansion period with Ag-scaffolds or free-peptide. B) The percentage of antigen-specific CD8+ T cells after 14 days of expansion as determined by antigen-tetramers and flow cytometry. C) The number of antigen-specific cells after 14 days of expansion in cultures expanded with Ag-scaffolds or free peptide. D) The fold expansion of antigen-specific cells after 14 days of expansion with either Ag-scaffolds or free-peptide.

**Supplementary Figure 3:**
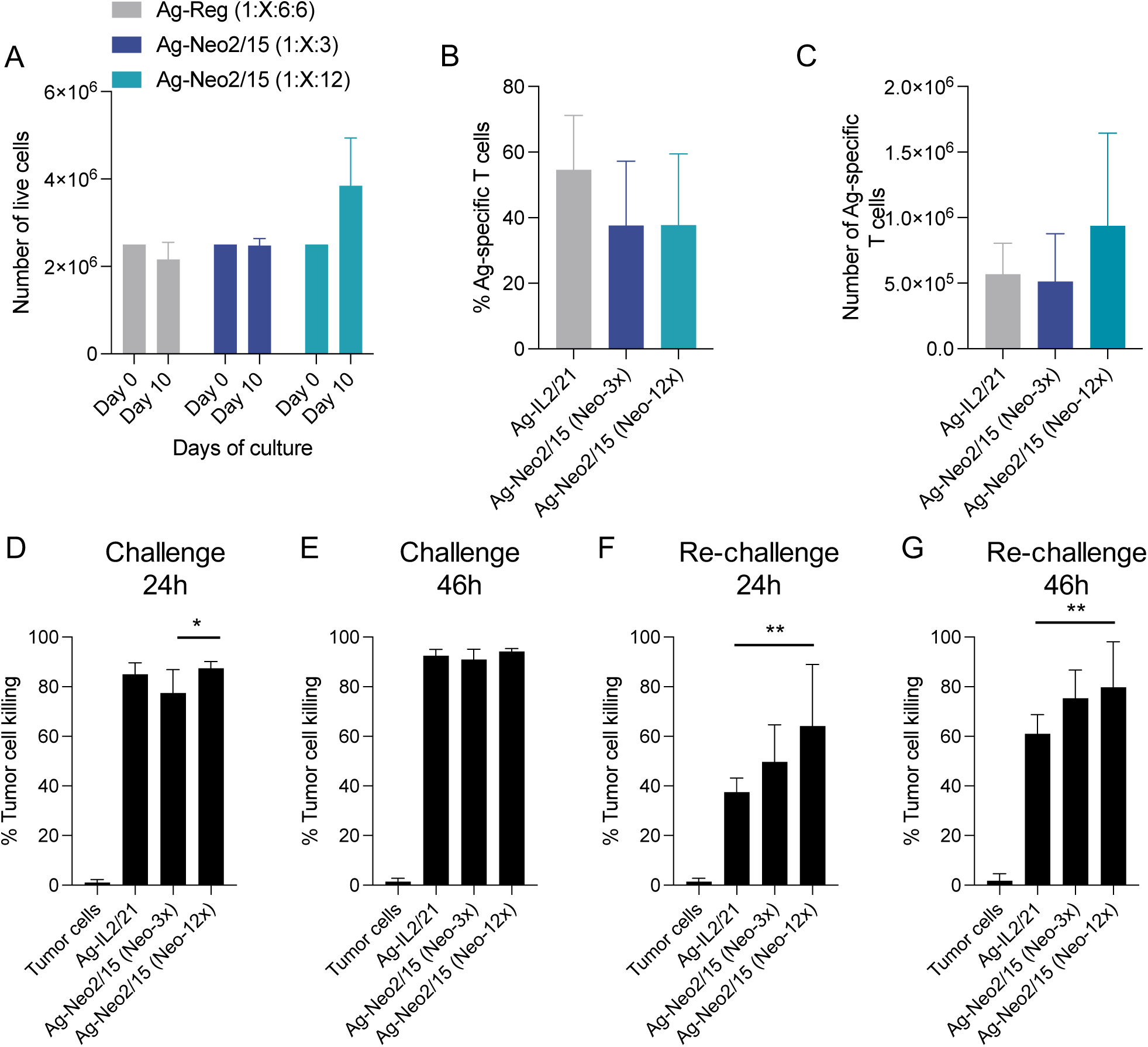
The expansion kinetics and cytotoxic capacity of CMV specific T cells expanded with Ag-Reg or Ag-Neo2/15 scaffolds. A) The number of live cells in cultures expanded with Ag-scaffolds for 10 days. B) % of antigen-specific T cells in cultures at day 0 and 10 as determined by tetramer-staining and flow cytometry. C) The number of antigen-specific T cells at day 0 and 10 of culture. The maximum target cell killing at D) 24h and E) 46h after initial co-cultivation with Ag-scaffold expanded T cells. The maximum cell killing at F) 24h and G) 46h after T cells are re-challenged with target cells.

**Supplementary Figure 4:**
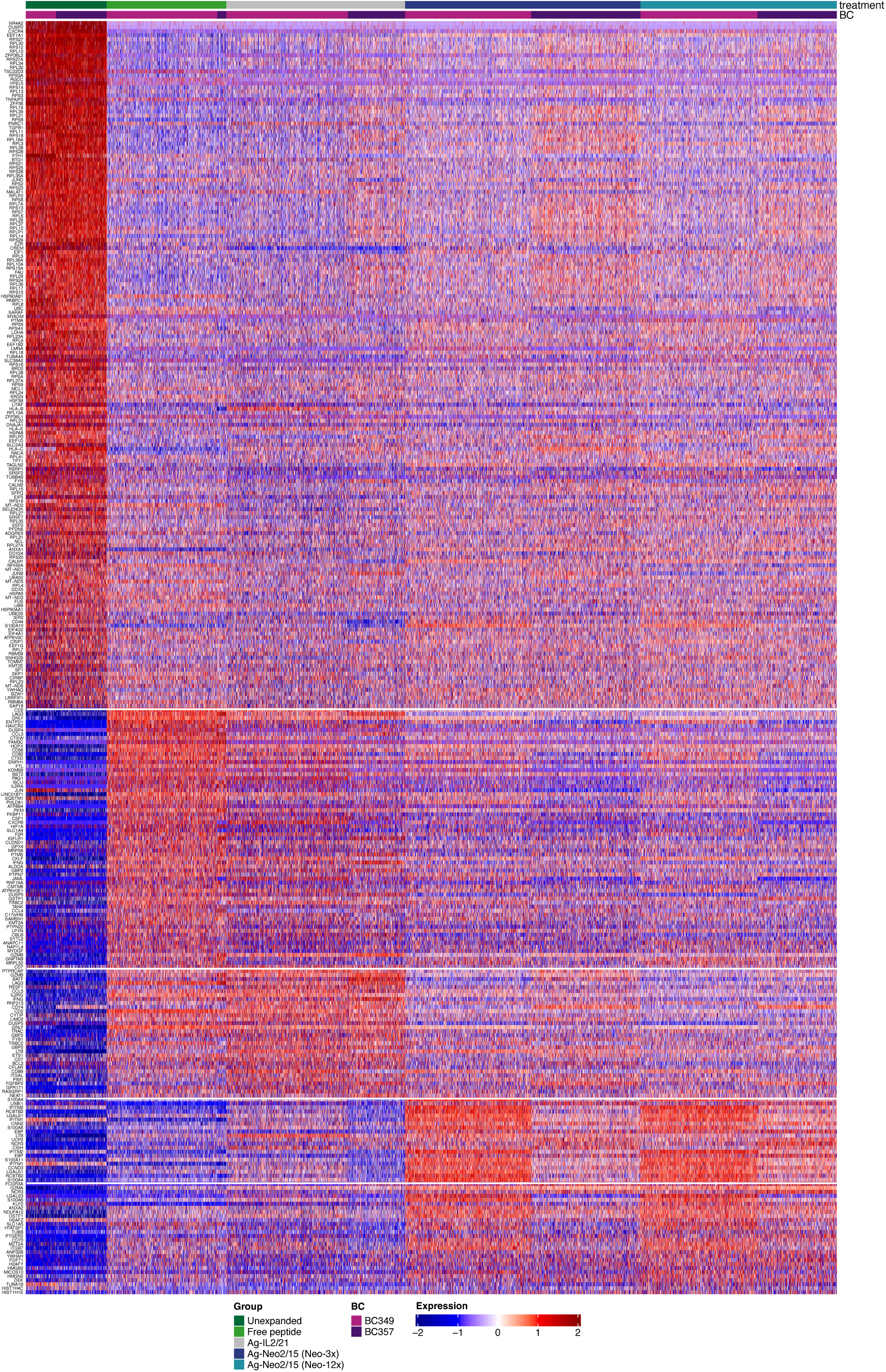
Heatmap of most differentially expressed genes for each expansion strategy. Heatmap from the single-cell analysis from the RNA panel, showing the most differentially expressed genes for each expansion strategy, clustering the cells fore each healthy donor within each expansion strategy.

**Supplementary Figure 5:**
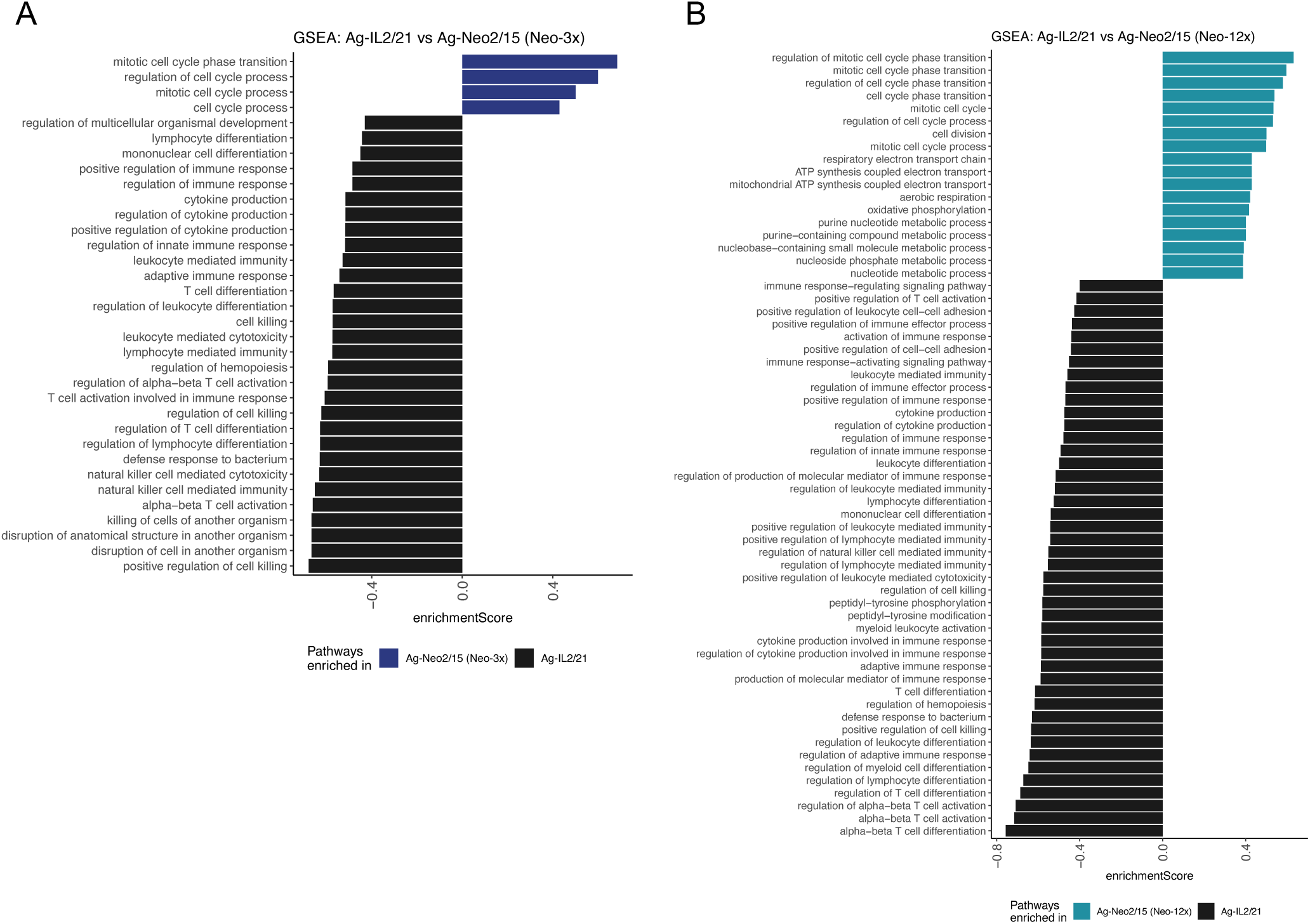
Summary of all the biological pathways found in the GSEA. Enrichment scores for the biological pathways, determined by the gene set enrichment analysis (GSEA) when comparing A) Ag-IL2/21 and Ag-Neo2/15 (Neo-3x) and when comparing B) Ag-IL2/21 and Ag-Neo2/15 (Neo-12x).

**Supplementary Figure 6:**
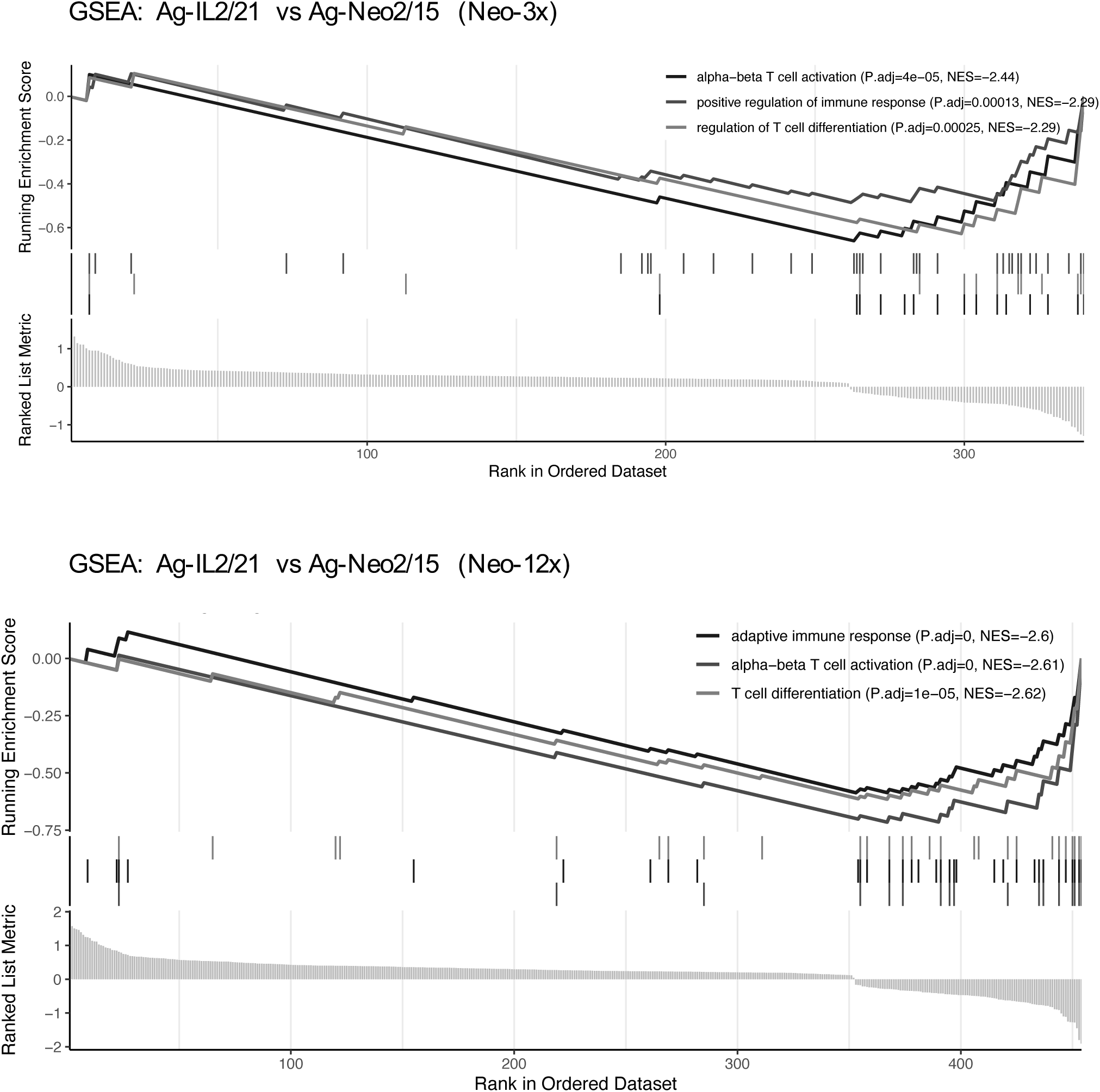
Selected biological pathways found in the GSEA enriched in Ag-IL2/21. Selected biological pathways determined by the gene set enrichment analysis when comparing Ag-IL2/21 vs Ag-Neo2/15 (Neo-3x) (top) and when comparing Ag-IL2/21 vs Ag-Neo2/15 (Neo-12x) (bottom).

**Supplementary Figure 7:**
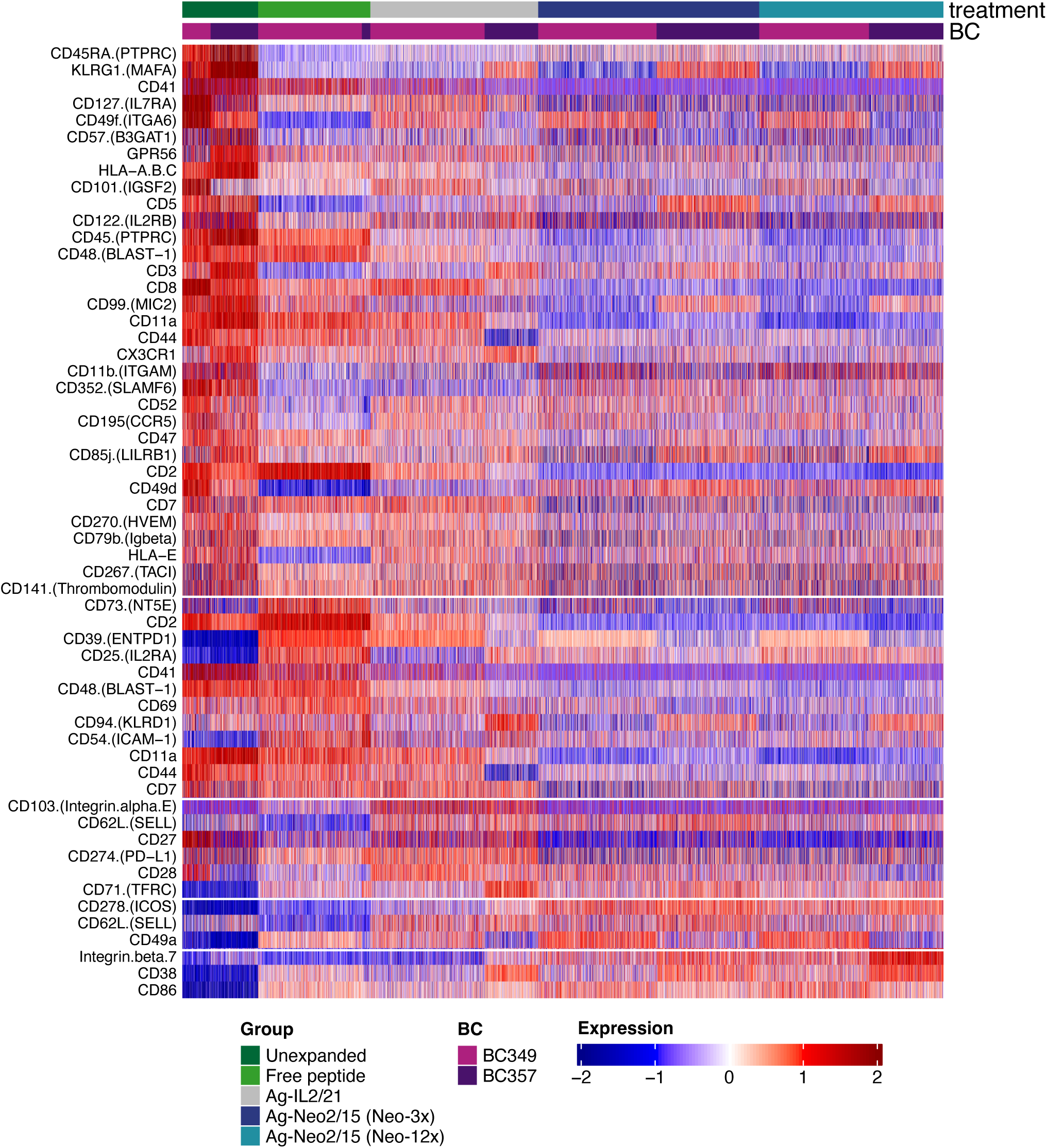
Most differentially expressed surface markers from the phenotype panel. Heatmap from the single-cell analysis phenotype panel, showing the most differentially expressed surface markers for each expansion strategy, clustering the cells fore each healthy donor within each expansion strategy.

**Supplementary Figure 8:**
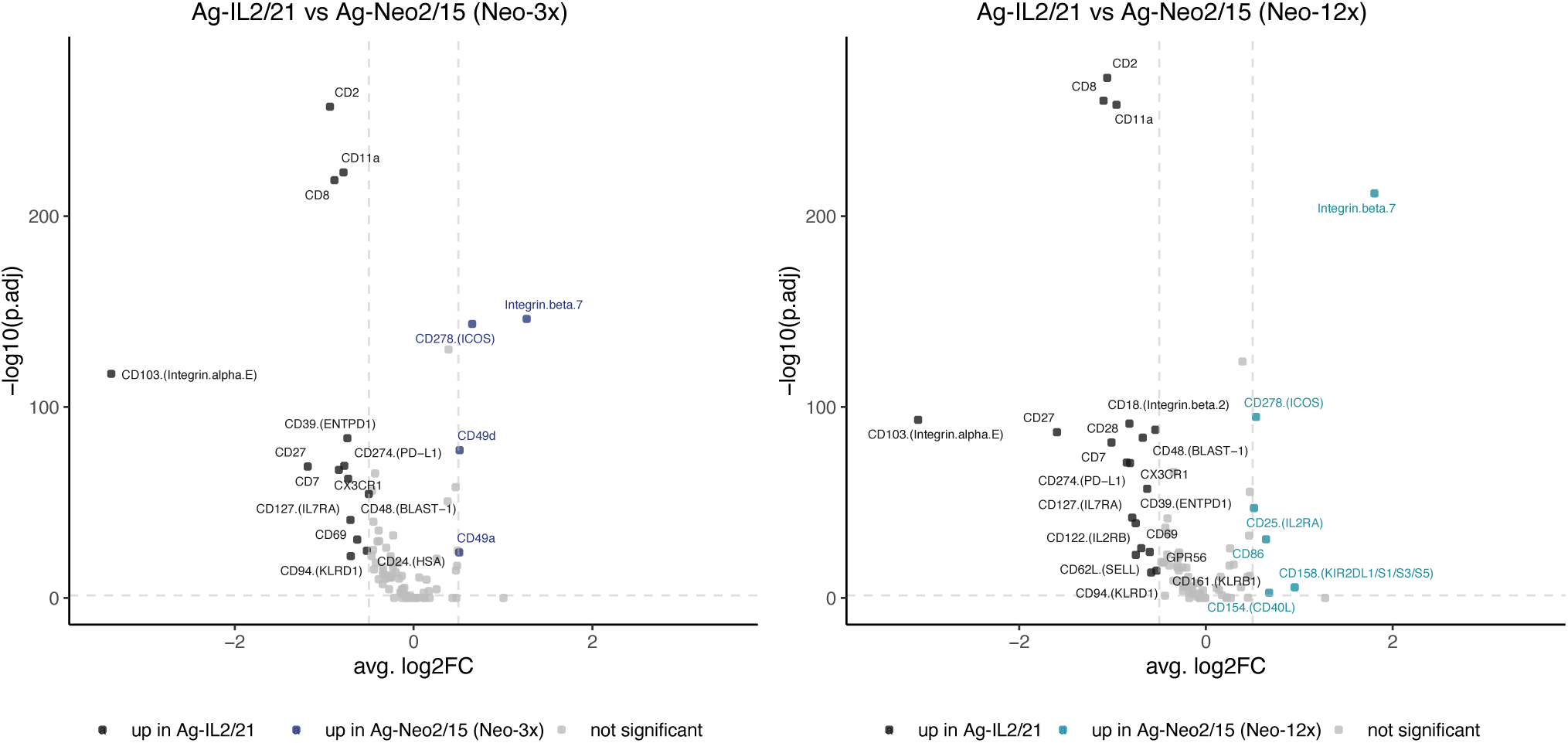
Most differentially expressed surface markers from the phenotype panel Volcano plot showing key differentially expressed surface markers from the phenotype panel, between the Ag-IL2/21 and Ag-Neo2/15 (Neo-3x) (left panel) and between the Ag-IL2/21 and Ag-Neo2/15 (Neo-12x) (right panel).

**Supplementary Figure 9:**
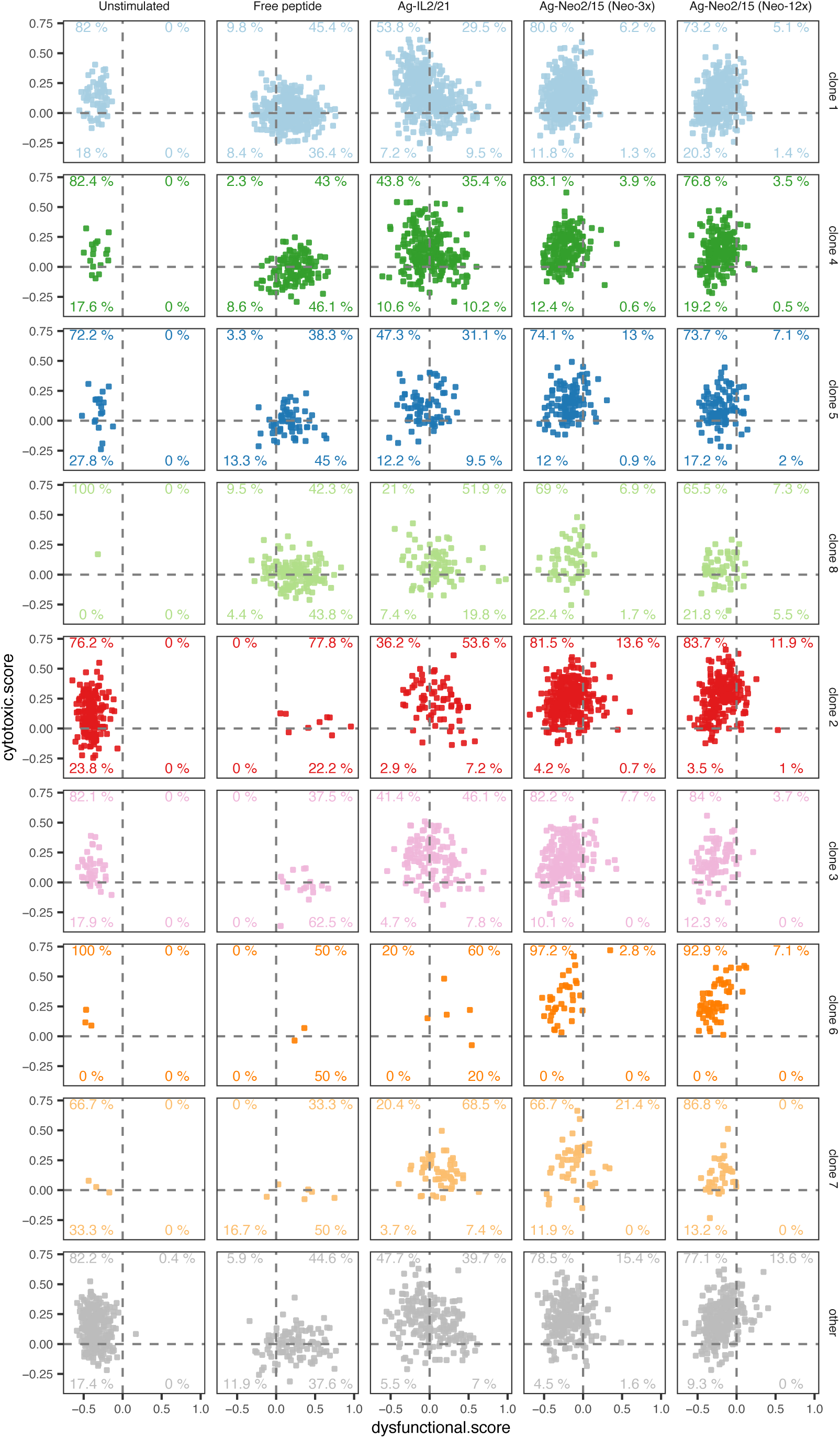
Relationship between the cytotoxic and dysfunctional score for the top four TCR clones for each donor. Quadrant scatter plots showing the relationship between the cytotoxic and dysfunctional score for each the top four TCR clones within each donor. Each scatter plot is divided into four quadrants. Quadrants with cyt+dys+ show cells where both scores are above the zero, cyt-dys-means that both scores are below zero, cyt+dys-represents cells where the cytotoxic score is above zero and the dysfunctional score is below zero and the cyt-dys+ are cells where the cytotoxic score is below zero and the dysfunctional score is above zero.

**Supplementary Table 1:**
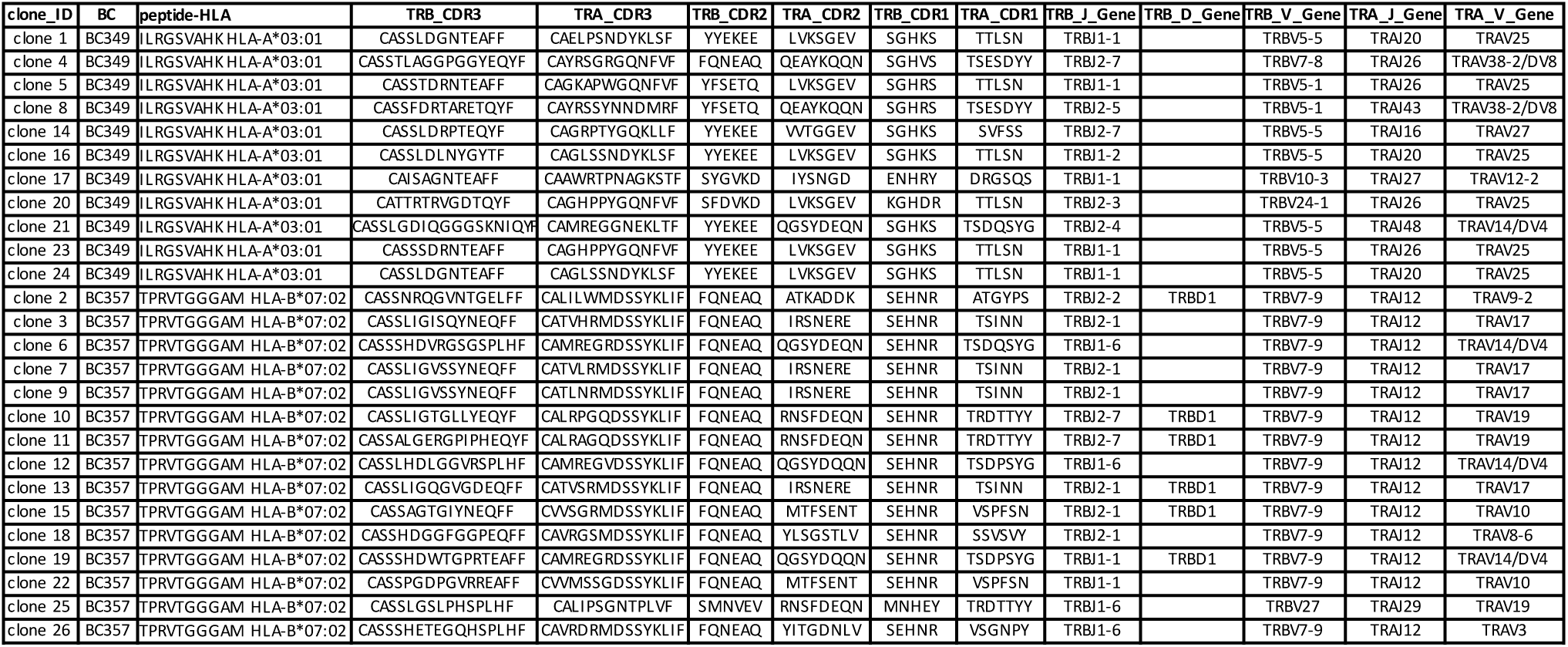
TCR sequences of ILR- and TPR-specific CD8+ T cells.

